# Age-Related Central Gain with Degraded Neural Synchrony in the Auditory Brainstem of Mice and Humans

**DOI:** 10.1101/2022.02.23.481643

**Authors:** Jeffrey A. Rumschlag, Carolyn M. McClaskey, James W. Dias, Lilyana B. Kerouac, Kenyaria V. Noble, Clarisse Panganiban, Hainan Lang, Kelly C. Harris

**Author notes:** Corresponding Author: Jeffrey A. Rumschlag.

## Abstract

Aging is associated with auditory nerve (AN) functional deficits and decreased inhibition in the central auditory system, amplifying central responses in a process known as central gain. Although central gain enhances response amplitudes, central gain may not restore disrupted response timing. In this translational study, we measured responses from the AN and auditory midbrain in younger and older mice and humans. We hypothesized that older mice and humans exhibit central gain without an improvement in inter-trial synchrony in the midbrain. Our data demonstrated greater age-related deficits in AN response amplitudes than auditory midbrain response amplitudes, as shown by significant interactions between neural generator and age group, indicating central gain in auditory midbrain. However, synchrony decreases with age in both the AN and midbrain responses. These results reveal age-related central gain without concomitant improvements in synchrony, consistent with those predictions based on decreases in inhibition. Persistent decreases in synchrony may contribute to auditory processing deficits in older mice and humans.

## 1. Introduction

Age-related loss of afferent AN fibers is well established, and this loss predicts auditory processing and speech recognition difficulties. However, these AN deficits appear to be ameliorated by central gain mechanisms, where the central nervous system compensates for a loss of afferent input by amplifying central responses (Dias et al., 2018; Gmehlin et al., 2011; Harris et al., 2012; Price et al., 2017; Tremblay et al., 2004; Woods and Clayworth, 1986; Pfefferbaum et al., 1979). Despite restoration of response amplitudes, auditory processing difficulties persist. One potential explanation for these continued difficulties is a disruption in neural synchrony. Neural synchrony is fundamental to auditory processing in difficult listening conditions, and increased neural jitter has been hypothesized to contribute to deficits observed in older adults. The current manuscript tests the hypothesis that while central gain may restore response amplitudes, deficits in neural synchrony persist and are propagated through the central auditory system.

Central gain is hypothesized to be associated with age-related declines in central inhibition. Aging is associated with declines in inhibitory transmission throughout the brain, possibly as a result of changes to gamma-aminobutyric acid (GABA) and glycine receptor composition (Caspary, 2008). Furthermore, the expression of markers of inhibitory interneurons decreases with age in mice (Brewton et al., 2016; Rogalla & Hildebrandt, 2020; Ueno et al., 2018), rats (Cisneros-Franco et al., 2018; Ouda et al., 2008; Ouellet & de Villers-Sidani, 2014), and humans (Mohan et al., 2018). The aging central auditory system may compensate for decreased afferent input with a reduction of inhibitory activity, resulting in the amplification of auditory signaling afferent to the AN, in a process known as central gain (Caspary et al., 2008).

To evaluate auditory central gain in younger and older mice and humans, we recorded compound action potentials (CAP) and auditory brainstem responses (ABR). The relationship between AN (ABR wave I or CAP N1) and midbrain (ABR wave V) responses can be used to estimate central gain. Central gain in the aging auditory system has been demonstrated by an increase in wave V/I ratio in animal models (Cai et al., 2018; Möhrle et al., 2016; Parthasarathy & Kujawa, 2018; Sergeyenko et al., 2013), and in humans (Grose et al., 2019; Psatta & Matei, 1988; Sand, 1991). This increase in wave V/I ratio arises either from larger response amplitudes at the midbrain (wave V) relative to AN or from an age-related decrease in ABR wave I amplitude without a decrease in wave V amplitude. Combining wave I and wave V into a single metric entangles their variabilities, limiting our ability to identify how different parts of the auditory system change with age, so this study instead uses linear mixed-effects regression (LMER) models to assess central gain. In this approach, central gain is indicated by an interaction between wave (wave I or wave V) and age group.

While central gain following acute insults has been well-characterized in animal models, less is known regarding chronic conditions like aging. Mouse models of acute cochlear insults, such as ouabain toxicity and noise-induced hearing loss, show that central gain manifests following a loss of afferent input. Eliminating >95% of Type-I SGN synapses and neuron somas via application of ouabain to the round window leads to a severe reduction in ABR wave I amplitude and a reduction in auditory-evoked activity in the inferior colliculus (Lang et al., 2010; Chambers et al., 2016). After 30 days, however, although ABR wave I amplitudes do not appear to recover significantly, midbrain responses are partially recovered. Central gain also develops following damaging broadband noise exposure, but peripheral neural and behavioral auditory deficits persist, as evidenced by decreased suprathreshold amplitudes and poor high frequency tone detection (Schrode et al., 2018). Similarly, age-related central gain does not appear to fully rescue behavioral auditory function.

In older adults, increased jitter in response timing is hypothesized to contribute to auditory processing deficits. Neural synchrony across trials, measured as the phase-locking value (PLV), has been demonstrated to predict speech recognition (Harris et al., 2021). In this study, we calculated the PLV of CAP and ABR responses to estimate neural synchrony (Harris et al., 2018, 2021, McClaskey et al., 2020). Models of amplitude and synchrony were then compared to determine whether age-related central gain improves synchrony in mice and human.

We hypothesized that central gain would be apparent in the aging auditory system of both mouse and human, as indicated by preserved midbrain responses (ABR wave V amplitudes), in contrast to decreased AN responses (ABR wave I or CAP N1 amplitudes). Furthermore, we predicted that the synchrony of the signals measured from both the peripheral and central portions of the auditory system would be significantly lower in older subjects relative to younger, suggesting central gain-related increases in response amplitudes are not reflected in a preservation or enhancement of neural synchrony. This results from this study demonstrate that central gain occurs without improvements in neural synchrony, and it informs future translational studies exploring the consequences of midbrain neural dyssynchrony for auditory perception.

## 2. Materials and Methods

### Mice

All studies were performed in accordance with the guidelines of the Institutional Animal Care and Use Committee of the Medical University of South Carolina (MUSC). CBA/CaJ mice were originally purchased from The Jackson Laboratory (Bar Harbor, ME) and bred in the MUSC Animal Research Facility. The mice were housed in a vivarium with a 12h light/dark cycle and given standard lab chow and water *ad libitum*. Included in this study are 14 younger mice (mean age = 2.5 (SD 0.6) months; 8 females; 24 ears) and 9 older mice (mean age = 25.8 (SD 3.5) months; 5 females; 16 ears). ABRs elicited by 85 dB SPL tone pips at 5.6, 11.3, and 40 kHz, were recorded from all mice. The average ABR wave I thresholds at 11.3 kHz were 22.9 (SD 4.6) dB SPL for the younger mice and 55 (SD 11.3) dB SPL for the older mice (5 kHz: younger: 41.4 (SD 8.3) dB SPL; older: 67.3 (SD 10.1) dB SPL. 40 kHz: younger: 14.6 (SD 10.9) dB SPL; older: 57.3 (SD 18.6) dB SPL). Group averaged ABR wave I thresholds are shown in Figure 1A.

**Figure 1.**
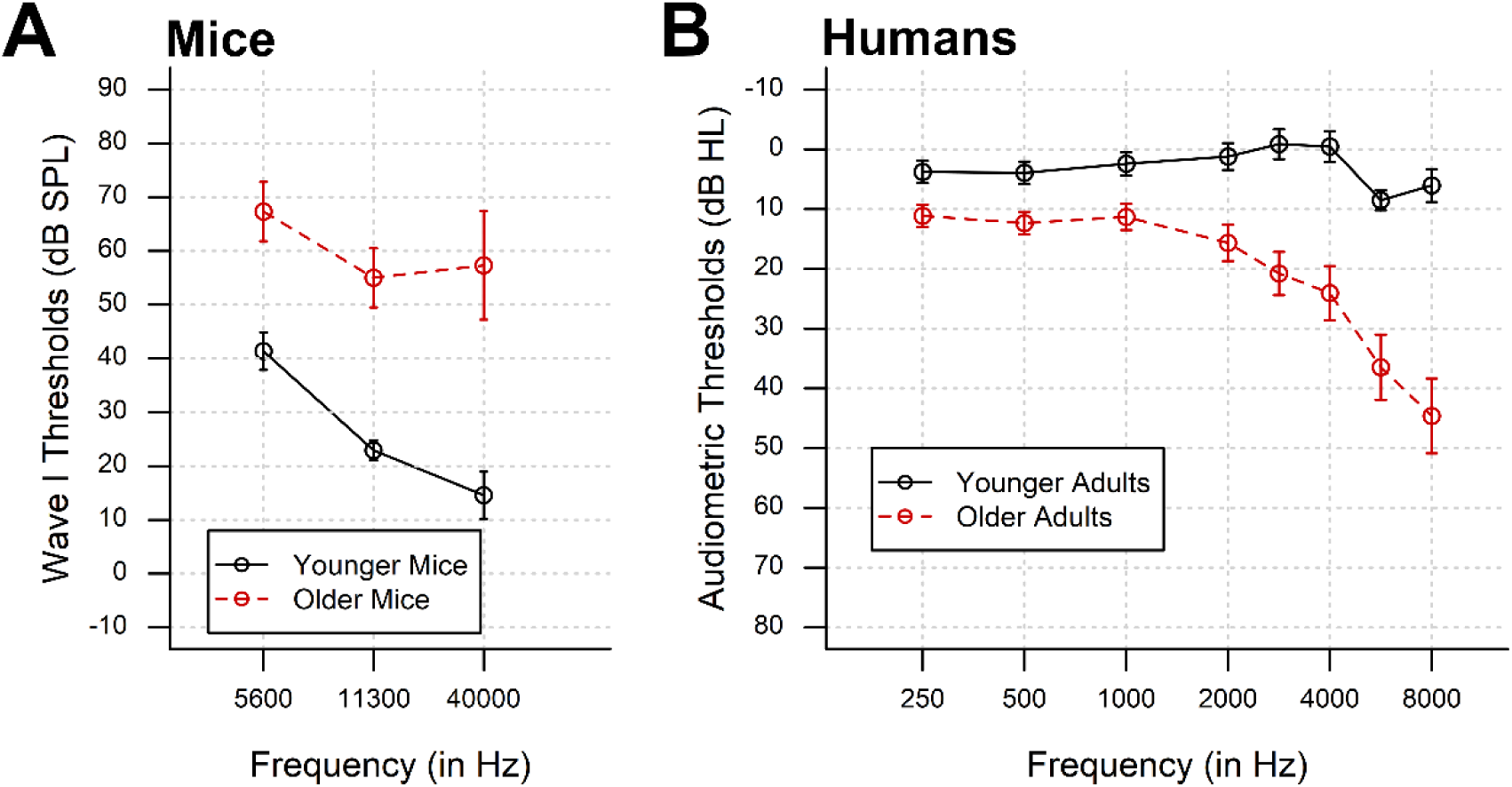
(A) ABR wave I thresholds in older mice are moderately elevated. (B) Right-ear pure-tone audiometric thresholds for younger and older human participants demonstrate greater hearing loss at higher frequencies. Error bars represent 95% confidence intervals.

### Human Participants

Participants included two groups of adults from the Charleston community: younger adults (N = 39; mean age = 24.1 (SD 3.1) years; 26 females) and older adults (N = 57; mean age = 66.0 (SD 6.6) years; 40 females). The participants were native English speakers with no otological or neurological impairments and had a Mini-Mental Status Examination score of at least 27. All recordings were done in the right ear. The younger participants had pure-tone thresholds ≤ 20 dB HL from 250 Hz to 8000 Hz. To examine central gain effects with age and age-related hearing loss, we included older adults with hearing thresholds ranging from normal limits to sloping or moderate-to-severe SNHL. Older adults were included if their hearing loss at or below 4 kHz did not exceed 65 dB HL. Group averages of audiometric thresholds with 95% confidence intervals are shown in Figure 1B. Participants provided written informed consent before participating in this study. Testing was initiated after approval by the Medical University of South Carolina’s Institutional Review Board.

### Mouse ABR Recordings

Mice were anesthetized via an intraperitoneal injection of a cocktail containing 20 mg/kg xylazine and 100 mg/kg ketamine. ABRs were recorded in an acoustically isolated booth. Subdermal needle electrodes were placed in the vertex of the scalp (recording electrode), in the ipsilateral mastoid (reference), and in the hind limb (ground). Electrodes were connected to a low-impedance head stage connected to a pre-amplifier (RA4LI/RA4PA, Tucker-Davis Technologies, Alachua, FL), and impedances were tested prior to each recording session and did not exceed 3 kΩ. The pre-amplifier was connected via optical cable to an RZ6 input/output device (Tucker-Davis Technologies), which was used to produce stimuli via BioSigRZ software (Tucker-Davis Technologies). Responses were recorded with RPvdsEx software (Tucker-Davis Technologies) at 24.414 kHz. ABRs were elicited by 85 dB SPL tone pips at 5.6 kHz, 11.3 kHz, and 40 kHz, which are represented in the apical, middle, and basal portions of the mouse cochlea, respectively. At least 515 tone pips of each frequency were presented. Closed-field stimuli were presented through an MF1 speaker (Tucker-Davis Technologies), coupled to a 3-mm diameter plastic tube and earpiece (total length, 1.6-1.8 cm), inserted into the ear canal. Calibration was performed using a 378C01 ICP microphone system (PCB Piezotronics, Inc., NY, USA), including a ¼” PCB 426 B03 032090 transducer (ICP@Sensor) and a model 480C02 battery-powered signal conditioner.

For each trace, if fewer than 300 trials were valid due to movement or noise artifacts, then that trace was excluded from further processing. Final analyses included recordings from 14 younger (5.6 kHz: 22 ears, 11.3 kHz: 24 ears, and 40 kHz: 24 ears) and 9 older mice (5.6 kHz: 11 ears, 11.3 kHz: 16 ears, and 40 kHz: 13 ears).

### Human CAP and ABR Recordings

CAPs and ABRs were recorded simultaneously in humans. Responses were elicited by 110 dB SPL, 100 μs rectangular pulses with alternating polarity, presented at 11.1 Hz to the right ear through an insert earphone (ER-3c; Etymotic Technologies), and responses were recorded from the right ear using a tympanic membrane electrode (Sanibel Supply, Eden Prairie, MN). The recording electrode was referenced to the left mastoid and grounded to an electrode placed on the low forehead, which was shared between the CAP and ABR setups. ABRs were recorded using an active electrode placed on the high forehead, referenced to the right mastoid, and grounded to the low forehead. The electrodes were connected to a custom headstage (Tucker Davis Technologies), which was connected to the bipolar channels of a Neuroscan SynAmpsRT amplifier in AC mode with 2010x gain (Compumedics USA, Charlotte, NC). Responses were recorded in blocks of 1100 trials (550 of each polarity) in CURRY (versions 7 and 8, Compumedics USA, Charlotte, NC) at a 20 kHz sampling rate and stored offline. During the recording, participants reclined in a chair in an acoustically and electrically shielded room. Participants were encouraged to sleep or rest quietly for the duration of the recording and to limit unnecessary or excessive movement.

### Peak Measurement

Recordings of continuous neural activity from mice and humans were analyzed in MATLAB (MathWorks, Natick, MA) using standard functions from EEGLAB (Delorme & Makeig, 2004) and ERPLAB (Lopez-Calderon & Luck, 2014). Recordings were bandpass filtered between 150 and 3000 Hz. The filtered recordings were epoched from -2 to 10 ms, relative to stimulus triggers, and baseline corrected to a pre-stimulus baseline of -2 ms to 0 ms (McClaskey et al., 2018). Aberrant individual responses were rejected based on a threshold of +/- 45 μV and subsequent visual inspection. The validated epochs were averaged, and the relevant peaks were identified: ABR wave I and V were identified in mouse recordings (Figure 2A), and CAP N1 and ABR wave V were identified in human recordings (Figure 2B). Peak selection was performed by at least two independent reviewers and assessed for repeatability across multiple runs. The reviewers were blinded to participant age group. Peak latencies and amplitudes were recorded for later analysis.

**Figure 2.**
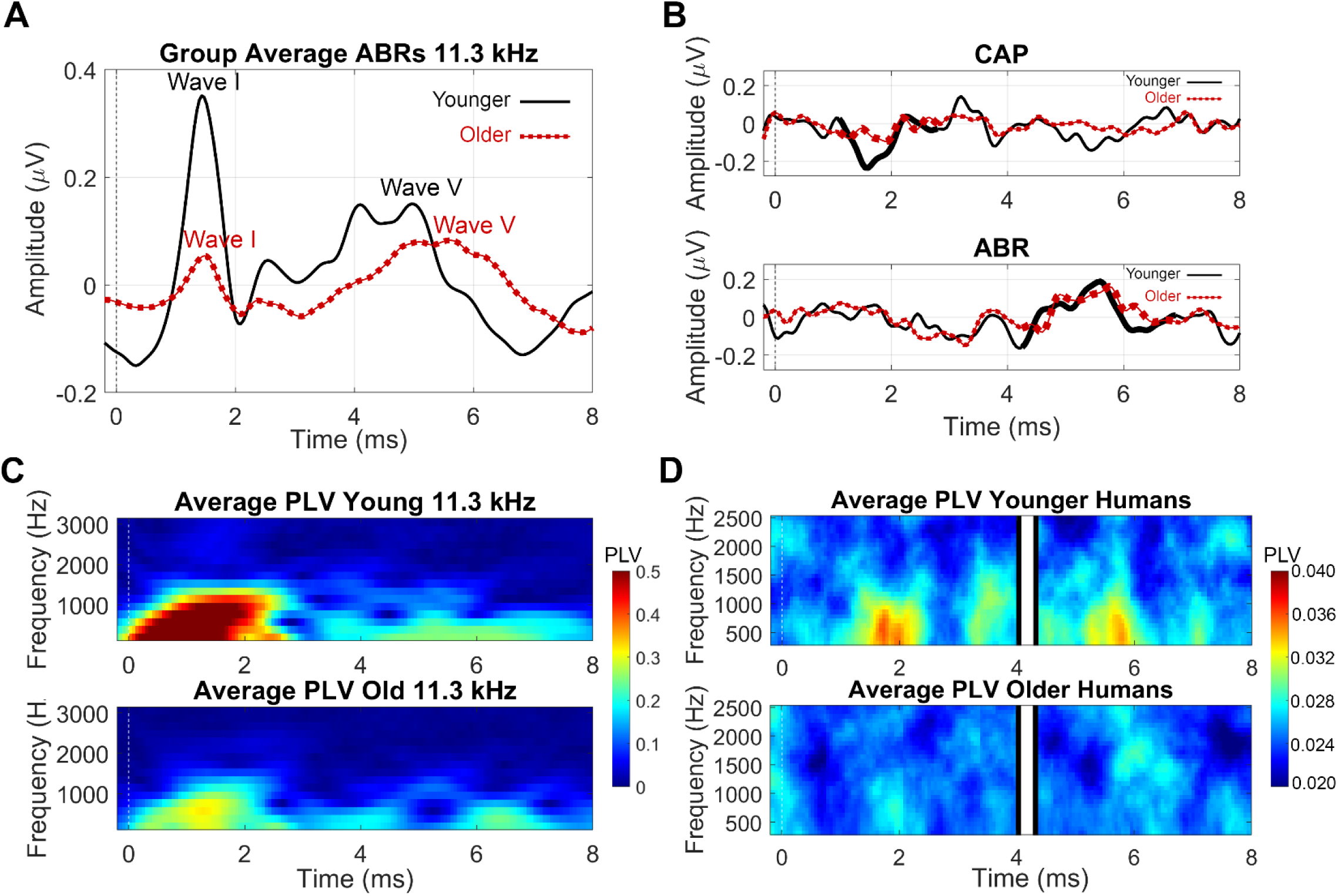
(A) Peak measurement locations for wave I and wave V are shown with vertical dotted lines on grand average ABR waveforms from younger (black line) and older (red line) mice, elicited by 11 kHz tone pips. (B) Average CAP and ABR waveforms recorded from younger (black line) and older (red line) humans are shown. The CAP N1 and ABR wave V are traced in thicker lines. (C) Average PLV heatmaps from mice depict synchronous activity corresponding to waves I and V of the ABR. (D) Average PLV heatmaps from human CAP (left panel) and ABR (right panel) recordings are shown on the same time axis, with 0-4 ms of the CAP and 4.3-8 ms of the ABR, to capture the CAP N1 and the ABR wave V.

### Synchrony (PLV)

PLV is a measure of the inter-trial coherence of the response, calculated for each time-frequency point as the magnitude of the trial-averaged phase vector. Whereas response amplitude is determined both by temporal jitter and by the response amplitudes within each trial, PLV reflects temporal jitter, independent from response amplitudes. Therefore, while PLV and response amplitudes both increase as stimulus level increases, as shown previously (Harris et al., 2018, 2021; McClaskey et al., 2020), examining these two measures together provides a means to dissociate synchrony from amplitude. PLV is calculated from a complex time-frequency transform, using the following equation, in which *F*_*k*_ (f, t) is the spectral estimate of trial *k* at frequency *f* and time *t*:

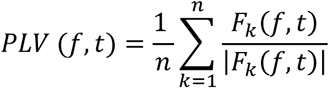

In this study, Hanning FFT tapers were applied to the continuous neural activity data, using the newtimef() function in EEGLAB. We analyzed 16 linearly spaced frequencies from 625 Hz to 2500 Hz, using a pad-ratio of 2 and a window size of 32 samples. For each of the peaks, the maximum PLV was extracted from a 2-ms window centered on the response latency (Figure 2C-D).

### Analytical Approach

In this study, central gain in humans was measured from CAP N1 and ABR wave V amplitudes. To allow for the comparison of these two measures while preserving the variability within each measure, standardized LMER models were used instead of amplitude ratios. We used LMER models to test for amplitude differences between wave I and wave V and between younger and older mice and human subjects. LMER is a non-parametric statistical approach that can test hypothesis-driven relationships between predictor and outcome variables while accounting for individual variability between subject groups (i.e., age-groups – younger and older) and variability between levels of a dependent variable that is nested within subjects (i.e., ABR waves – I and V). We employed a hierarchical model-testing approach to determine whether aging affects AN and midbrain responses differently. We fit hypothesis-driven LMER models to the AN and midbrain response measurements using the lme4 package for R (x64 v4.0.5). Amplitude and PLV at each frequently were modeled separately (e.g., Harris et al., 2018, 2021; McClaskey et al., 2020).

First, we tested the degree to which amplitude or PLV was different between age groups and between ABR wave I and wave V (in mice) or between CAP N1 and ABR wave V (in humans). To do this, age-group and wave were added to a main-effects LMER model with measure (amplitude or PLV) as the outcome variable and individual (human or mouse) as a random factor. If central gain is present in the aging auditory system, then ABR wave I or CAP N1 magnitude will decrease more than ABR wave V amplitude with age. To test this hypothesis, we added the wave number and age group interaction term to the main-effects model to determine if model fit was improved. If aging differentially affects the AN and midbrain measurements, as would be expected with central gain, then the interaction will be significant, and including the interaction term will improve model fit. Post-hoc linear models (LMs) were conducted to explore significant interactions. For mice, these models were tested for each test frequency separately. To determine whether our results could be accounted for by age-related hearing loss or sex differences, a measure of audiometric threshold (ABR wave I threshold for mice; pure-tone average from 250 Hz to 8000 Hz for humans) and individual sex were added to the LMER models to determine if model fit was improved.

## 3. Results

### 3.1 ABR amplitudes demonstrate central gain in aging mice

Figure 3 shows boxplots representing the response amplitudes for each of the mouse age groups for waves I and V in response to the tone pip stimuli of 5.6 kHz (Fig. 3A), 11.3 kHz (Fig. 3B), and 40 kHz (Fig. 3C). Including the interaction term of wave number and age group improved model fit for ABR response amplitudes over the main effects model for 5.6 kHz (χ^2^(1) = 22.371, p < 0.001), 11.3 kHz (χ^2^(1) = 23.671, p < 0.001), and 40 kHz (χ^2^(1) = 11.193, p < 0.001), showing that age differentially impacts the AN and midbrain. The parameters for these models and post-hoc LM tests are reported in Table 1. Wave I amplitudes were larger than wave V amplitudes and the amplitudes of both waves decreased with age for all test frequencies. Post-hoc tests exploring the significant interaction of age group and wave for each test frequency (Table 1) found that age-related amplitude deficits in the AN exceeded those observed in the midbrain (i.e., larger βs; see also Fig. 3). Adding hearing thresholds or sex did not improve the fit of any of these models, nor did they prove to be significant predictors of response amplitudes.

**Figure 3.**
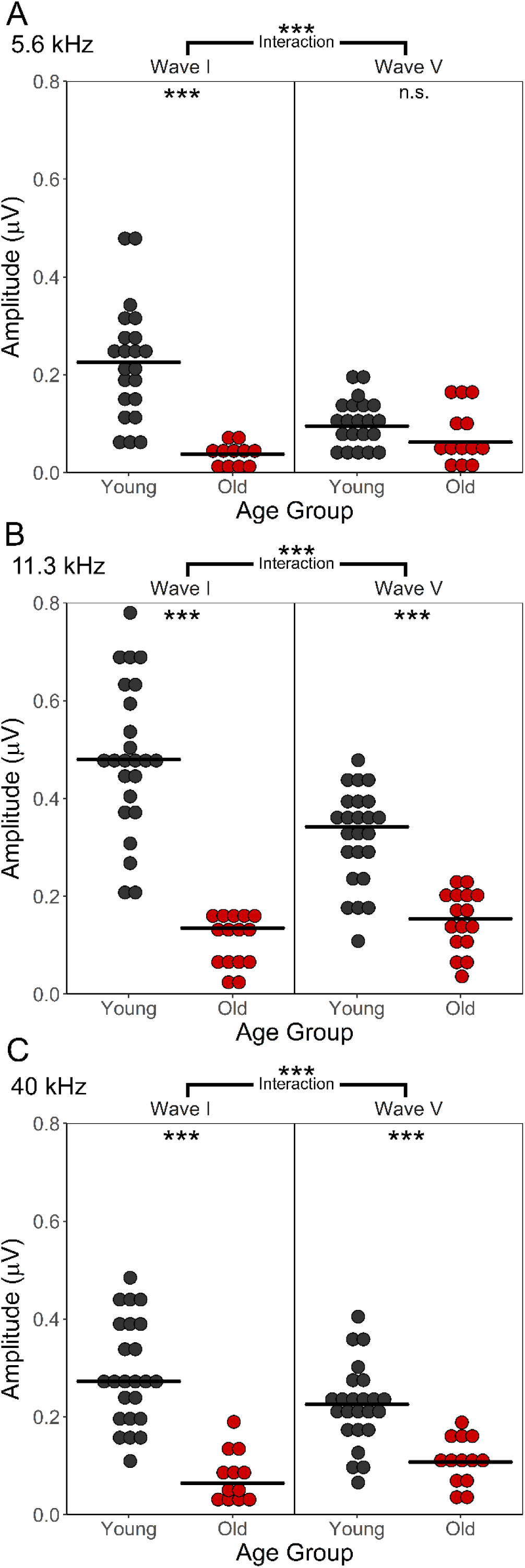
ABR wave I and wave V amplitudes from younger and older CBA/CaJ mice in response to 5.6 kHz **(A)**, 11.3 kHz **(B)**, and 40 kHz **(C)** tone pips demonstrate age-related central gain. Asterisks on brackets spanning wave I and wave V indicate a significant interaction of wave and age group. Asterisks within the age comparison plots indicate a significant post-hoc age-related effect. Detailed statistical results from these analyses appear in **Table 1**. *n*.*s. not significant; *p<0*.*05; **p<0*.*01; ***p<0*.*001*.

**Table 1.** Mouse ABR Amplitude

| | $B$ | $SE_B$ | $\beta$ | $SE_\beta$ | $t$ | $p$ | |
| --- | --- | --- | --- | --- | --- | --- | --- |
| ***Mouse 5.6 kHz Amplitude: Age and Wave Interaction Model*** |  |  |  |  |  |  |  |
| Intercept | 0.226 | 0.017 | 0.291 | 0.123 | 13.552 | <0.001 | *** |
| Age Group | -0.201 | 0.027 | -0.903 | 0.123 | -7.315 | <0.001 | *** |
| Wave | -0.129 | 0.020 | -0.582 | 0.148 | -6.433 | <0.001 | *** |
| Age Group x Wave | 0.179 | 0.033 | 0.806 | 0.149 | 5.427 | <0.001 | *** |
| Post-hoc Linear Models |  |  |  |  |  |  |  |
| **Wave I** |  |  |  |  |  |  |  |
| Intercept | 0.226 | 0.020 | 0.000 | 0.119 | 11.072 | <0.001 | *** |
| Age Group | -0.201 | 0.034 | -0.721 | 0.121 | -5.976 | <0.001 | *** |
| **Wave V** |  |  |  |  |  |  |  |
| Intercept | 0.097 | 0.012 | 0.000 | 0.169 | 8.193 | <0.001 | *** |
| Age Group | -0.021 | 0.019 | -0.188 | -0.171 | -1.099 | 0.280 |  |
| ***Mouse 11.3 kHz Amplitude: Age and Wave Interaction Model*** |  |  |  |  |  |  |  |
| Intercept | 0.486 | 0.022 | 0.215 | 0.092 | 22.258 | <0.001 | *** |
| Age Group | -0.382 | 0.035 | -1.021 | 0.092 | -11.063 | <0.001 | *** |
| Wave | -0.162 | 0.024 | -0.429 | 0.100 | -6.834 | <0.001 | *** |
| Age Group x Wave | 0.208 | 0.038 | 0.557 | 0.100 | 5.538 | <0.001 | *** |
| Post-hoc Linear Models |  |  |  |  |  |  |  |
| **Wave I** |  |  |  |  |  |  |  |
| Intercept | 0.486 | 0.026 | 0.000 | 0.087 | 19.021 | <0.001 | *** |
| Age Group | -0.382 | 0.040 | -0.838 | 0.089 | -9.454 | <0.001 | *** |
| **Wave V** |  |  |  |  |  |  |  |
| Intercept | 0.323 | 0.017 | 0.000 | 0.112 | 18.661 | <0.001 | *** |
| Age Group | -0.174 | 0.027 | -0.717 | 0.113 | -6.339 | <0.001 | *** |
| ***Mouse 40 kHz Amplitude: Age and Wave Interaction Model*** |  |  |  |  |  |  |  |
| Intercept | 0.288 | 0.017 | 0.123 | 0.119 | 16.688 | <0.001 | *** |
| Age Group | -0.215 | 0.029 | -0.886 | 0.119 | -7.447 | <0.001 | *** |
| Wave | -0.065 | 0.017 | -0.246 | 0.119 | -3.746 | <0.001 | *** |
| Age Group x Wave | 0.102 | 0.029 | 0.421 | 0.120 | 3.515 | 0.001 | ** |
| Post-hoc Linear Models |  |  |  |  |  |  |  |
| **Wave I** |  |  |  |  |  |  |  |
| Intercept | 0.288 | 0.019 | 0.000 | 0.110 | 15.340 | <0.001 | *** |
| Age Group | -0.215 | 0.032 | -0.754 | 0.111 | -6.790 | <0.001 | *** |
| **Wave V** |  |  |  |  |  |  |  |
| Intercept | 0.224 | 0.015 | 0.000 | 0.133 | 14.993 | <0.001 | *** |
| Age Group | -0.113 | 0.025 | -0.604 | -0.135 | -4.486 | <0.001 | *** |

Notes: \*p < 0.05, \*\*p < 0.01, \*\*\*p < 0.001.

### 3.2 ABR phase-locking values indicate degraded neural synchrony in aging mice

Figure 4 shows the wave I and wave V PLV for each age group at 5.6 kHz (Fig. 4A), 11.3 kHz (Fig. 4B), and 40 kHz (Fig. 4C). As reported in Table 2, across test-frequencies, PLV was smaller for older mice and for wave V. Including the interaction term for age group and wave improved model fit for the 5.6 kHz stimuli (Table 2, χ^2^(1) = 11.958, p < 0.001). Post-hoc linear models found that age-related deficits in neural synchrony were greater in the AN than the midbrain (larger β). The fit of our main effects model including age-group and wave was not improved by adding the interaction between wave and age group for the 11.3 kHz (χ^2^(1) = 0.377, p = 0.539) or 40 kHz (χ^2^(1) = 0.037, p = 0.848) test frequencies, suggesting that the synchrony of the responses to tone pips at these frequencies is uniformly degraded with age in both the peripheral and central portions of the auditory system. Adding hearing thresholds and sex did not improve the fit of any of these models, nor did they prove to be significant predictors of PLV.

**Figure 4.**
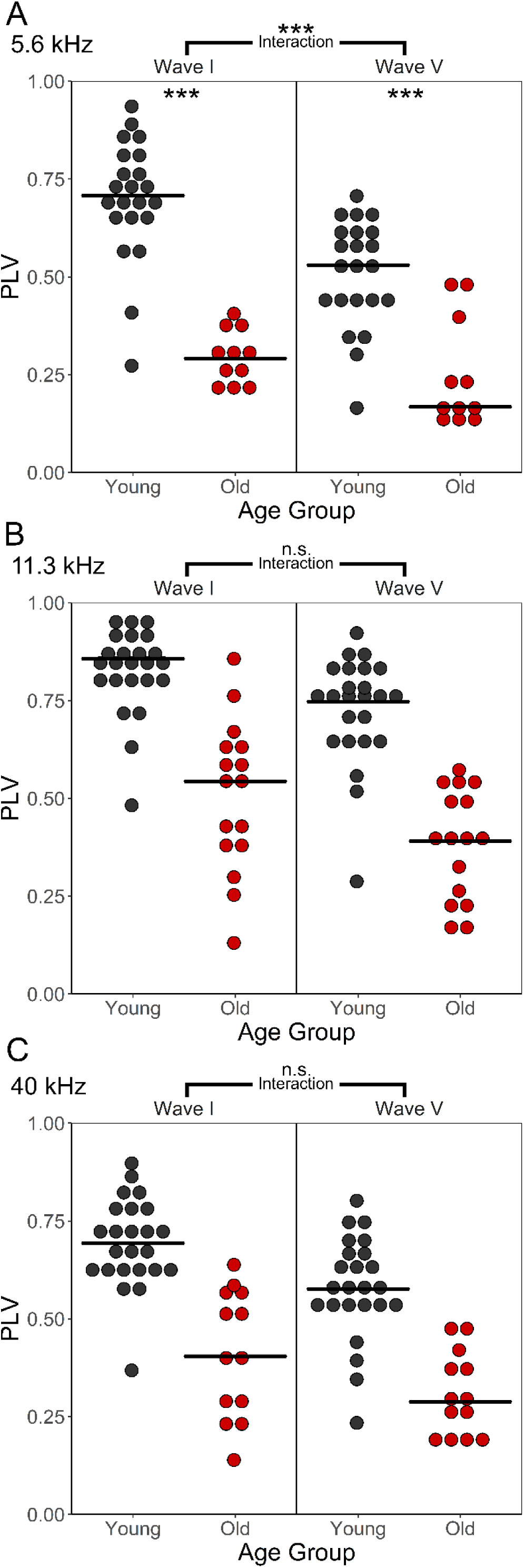
ABR wave I and wave V phase-locking values measured from younger and older CBA/CaJ mice in response to 5.6 kHz **(A)**, 11.3 kHz **(B)**, and 40 kHz **(C)** tone pips demonstrate age-related degradation of neural synchrony in peripheral and central portions of the auditory system, which appear to be less severe in the midbrain at lower frequencies. Neural synchrony measurements (PLV) for 5.6 kHz **(A)** show a significant interaction of age group and wave, whereas PLV for 11.3 kHz **(B)** and 40 kHz **(C)** do not. Asterisks on brackets spanning wave I and wave V indicate a significant interaction of wave and age group. Asterisks within age comparison plots indicate a significant post-hoc age-related effect. Detailed statistical results from these analyses appear in **Table 2**. *n*.*s. not significant; *p<0*.*05; **p<0*.*01; ***p<0*.*001*.

**Table 2.** Mouse PLV 5.6 kHz

| | $B$ | $SE_B$ | $\beta$ | $SE_\beta$ | $t$ | $p$ | |
| --- | --- | --- | --- | --- | --- | --- | --- |
| ***Mouse 5.6 kHz PLV: Age and Wave Interaction Model*** |  |  |  |  |  |  |  |
| Intercept | 0.700 | 0.029 | 0.300 | 0.103 | 24.401 | <0.001 | *** |
| Age Group | -0.407 | 0.047 | -0.895 | 0.104 | -8.650 | <0.001 | *** |
| Wave | -0.190 | 0.026 | -0.600 | 0.092 | -7.406 | <0.001 | *** |
| Age Group x Wave | 0.154 | 0.042 | 0.340 | 0.093 | 3.666 | <0.001 | *** |
| Post-hoc Linear Models |  |  |  |  |  |  |  |
| **Wave I** |  |  |  |  |  |  |  |
| Intercept | 0.700 | 0.028 | 0.000 | 0.093 | 25.288 | <0.001 | *** |
| Age Group | -0.407 | 0.045 | -0.842 | 0.094 | -8.965 | <0.001 | *** |
| **Wave V** |  |  |  |  |  |  |  |
| Intercept | 0.510 | 0.030 | 0.000 | 0.127 | 17.187 | <0.001 | *** |
| Age Group | -0.253 | 0.049 | -0.671 | 0.129 | -5.192 | <0.001 | *** |
| ***Mouse 11.3 kHz PLV: Age and Wave Model*** |  |  |  |  |  |  |  |
| Intercept | 0.832 | 0.028 | 0.247 | 0.101 | 29.530 | <0.001 | *** |
| Age Group | -0.332 | 0.042 | -0.736 | 0.094 | -7.842 | <0.001 | *** |
| Wave | -0.110 | 0.018 | -0.493 | 0.080 | -6.189 | <0.001 | *** |
| ***Mouse 40 kHz PLV: Age and Wave Model*** |  |  |  |  |  |  |  |
| Intercept | 0.692 | 0.025 | 0.292 | 0.111 | 27.921 | <0.001 | *** |
| Age Group | -0.276 | 0.036 | -0.694 | 0.091 | -7.672 | <0.001 | *** |
| Wave | -0.112 | 0.024 | -0.584 | 0.128 | -4.565 | <0.001 | *** |

Notes: \*p < 0.05, \*\*p < 0.01, \*\*\*p < 0.001.

### 3.3 AN and midbrain response amplitudes demonstrate central gain in older humans

CAP N1 and ABR wave V response amplitudes are summarized in Figure 5. Including the interaction term between wave and age group improved the fit of our main effects LMER model (χ^2^(1) = 4.272, p = 0.039). Table 3 summarizes the parameters of this interaction model. Wave V amplitudes were larger than CAP N1 amplitudes, and amplitudes across the CAP N1 and ABR wave V decreased with age (because CAP N1 is negative, β>0 means that the CAP N1 magnitude decreased), but age-group differences in amplitude differed between the CAP N1 and the ABR wave V. Post-hoc linear models found that the amplitude of the CAP N1 response is diminished in older adults, whereas the ABR wave V response is not significantly different between younger and older adults, suggesting central gain in the midbrain of older listeners. Adding hearing thresholds and sex did not improve the fit of this model, nor did they prove to be significant predictors of response amplitudes.

**Figure 5.**
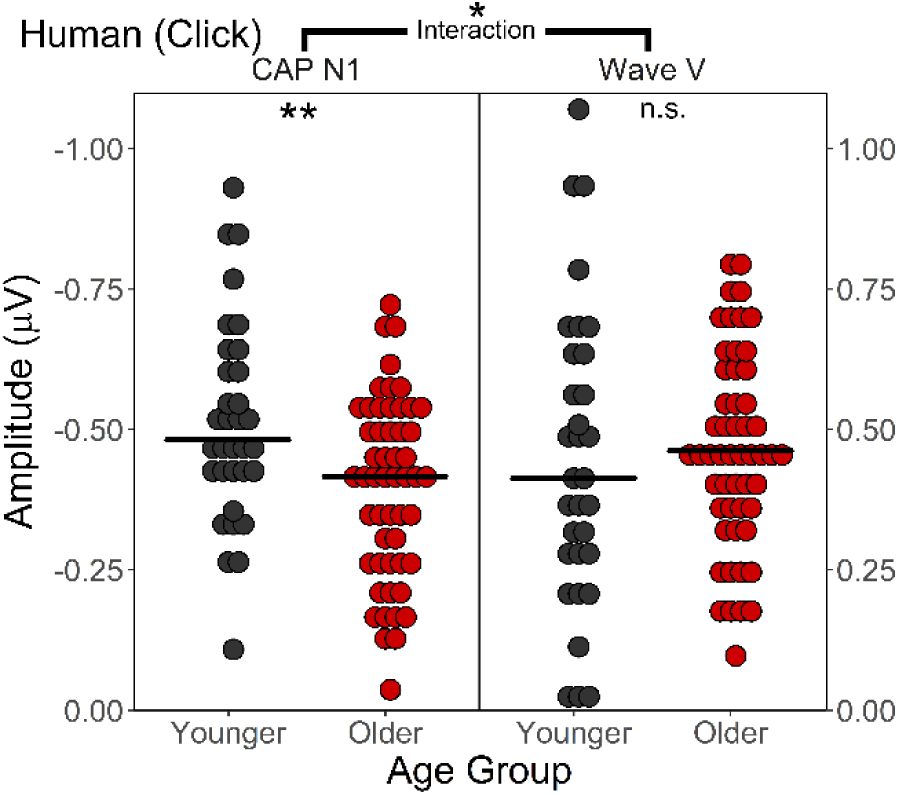
CAP N1 and ABR wave V amplitudes measured from younger and older human participants demonstrate age-related central gain. The asterisk on the bracket spanning CAP N1 amplitude and ABR wave V amplitude indicates a significant interaction of wave and age group, indicating that the amplitude of the AN response and the amplitude of the midbrain response are differentially affected by age. Asterisks within the age comparison plots indicate the significance of post-hoc age-related linear models. Detailed statistical results from these analyses appear in Table 3. *n*.*s. not significant; *p<0*.*05; **p<0*.*01; ***p<0*.*001*.

**Table 3.** Human CAP and ABR Amplitude (Click)

|  | *B* | *SE<sub>B</sub>* | *β* | *SE<sub>β</sub>* | *t* | *p* |  |
| --- | --- | --- | --- | --- | --- | --- | --- |
| *Human Amplitude: Age and Wave Interaction Model* |  |  |  |  |  |  |  |
| Intercept | -0.519 | 0.034 | -0.854 | 0.043 | -15.162 | <0.001 | *** |
| Age Group | 0.131† | 0.044 | 0.128† | 0.043 | 2.942 | 0.004 | ** |
| Wave | 0.984 | 0.050 | 1.799 | 0.063 | 19.607 | <0.001 | *** |
| Age Group x Wave | -0.133 | 0.065 | -0.130 | 0.063 | -2.056 | 0.041 | * |
| Post-hoc Linear Models |  |  |  |  |  |  |  |
| **CAP N1** |  |  |  |  |  |  |  |
| Intercept | -0.519 | 0.032 | 0.000 | 0.098 | -16.168 | <0.001 | *** |
| Age Group | 0.131† | 0.042 | 0.308† | 0.098 | 3.137 | 0.002 | ** |
| **ABR Wave V** |  |  |  |  |  |  |  |
| Intercept | 0.465 | 0.039 | 0.000 | 0.108 | 11.900 | <0.001 | *** |
| Age Group | -0.002 | 0.050 | -0.005 | 0.109 | -0.045 | 0.965 |  |

Notes: \*p < 0.05, \*\*p < 0.01, \*\*\*p < 0.001. †Because the CAP N1 amplitude is negative, these positive B and β values denote a decrease in response magnitude with age.

|  |  |  |  |  |  |  |  |
| --- | --- | --- | --- | --- | --- | --- | --- |
| *Human PLV: Age and Wave* |  |  |  |  |  |  |  |
| Intercept | 0.066 | 0.002 | 0.033 | 0.099 | 43.887 | <0.001 | *** |
| Age Group | -0.006 | 0.002 | -0.260 | 0.077 | -3.371 | 0.001 | ** |
| Wave | -0.001 | 0.001 | -0.075 | 0.132 | -0.570 | 0.570 |  |

Notes: \*p < 0.05, \*\*p < 0.01, \*\*\*p < 0.001.

### 3.4 Neural synchrony is degraded in human AN and auditory midbrain

CAP N1 and ABR wave V PLV are summarized in Figure 6. Including the interaction of age and wave in the model did not improve the fit of our main effects model, suggesting that PLV decreases with age similarly in the CAP N1 and ABR wave V. As reported in Table 3, PLV was smaller for older listeners and wave V PLV was smaller than CAP N1 PLV. Adding hearing thresholds and sex did not improve the fit of this model, nor did they prove to be significant predictors of PLV.

**Figure 6.**
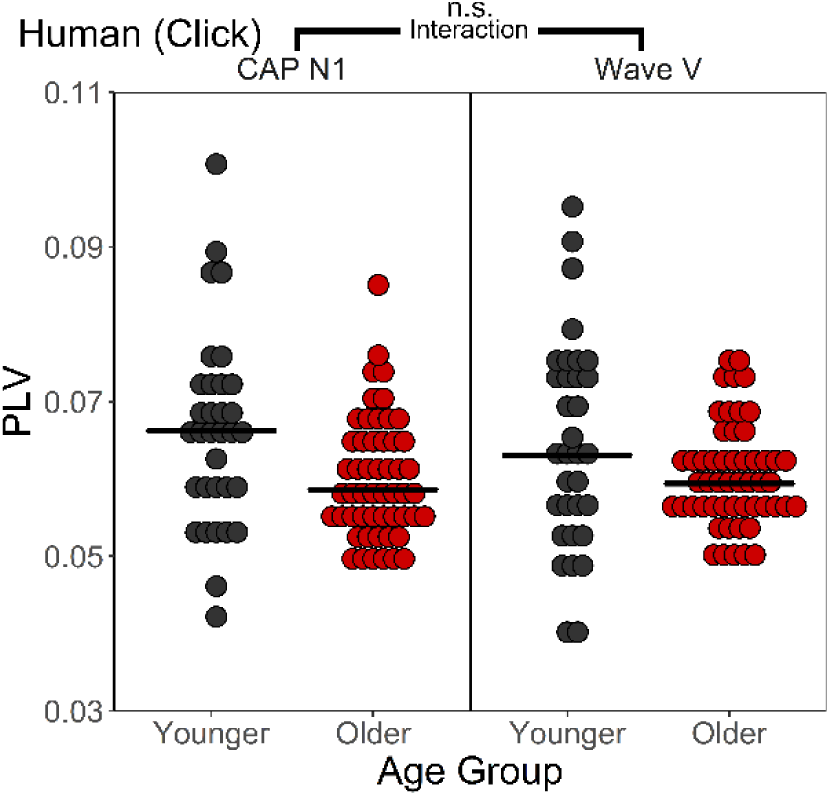
CAP N1 and ABR wave V phase-locking values measured from younger and older human participants demonstrate that degradation of neural synchrony occurs in both the peripheral and central portions of the aging auditory system. There is no significant interaction between age group and response source for CAP N1 PLV and ABR wave V PLV. The significant effect of age group (independent of response source) is indicated by asterisks in parentheses. Detailed statistical results from these analyses appear in Table 3. *n*.*s. not significant; *p<0*.*05; **p<0*.*01; ***p<0*.*001*

## 4. Discussion

Age-related loss and dysfunction of AN fibers are well-established. This loss co-occurs with a loss of neural synchrony, contributing to auditory processing deficits with age. However, the extent to which the central auditory system can compensate for these deficits is largely unknown. Our results suggest that age-related decreases in afferent input appear to be partially ameliorated by central gain in the brainstem in both mice and humans. Building upon these results, we provide evidence for the first time for persistent declines in neural synchrony. Taken together, restoration of response amplitudes with concomitant decreases in neural synchrony are consistent with animal models of decreased inhibition.

### Evidence of central gain in the aging auditory system

Amplitude-based analyses of ABR recordings from younger and older mice and humans demonstrate central gain in the aging mammalian auditory system. While the amplitudes of the responses generated in the AN are lower in older adults, the responses generated in the auditory midbrain are unaffected by age in humans (Figure 5) and significantly less affected by age in mice (Figure 3). These results are consistent with prior studies of the aging auditory system in mice (Parthasarathy & Kujawa, 2018; Sergeyenko et al., 2013), and in humans (Grose et al., 2019; Psatta & Matei, 1988; Sand, 1991). The model-testing approach taken in this study demonstrates that the patterns of amplitude and synchrony across regions are not dependent on hearing thresholds. The age-related decrease in suprathreshold ABR wave I and CAP N1 amplitudes have been attributed to a loss of low-spontaneous rate (SR) fibers (Schmiedt, 2010), age-related changes to myelination of the AN (Xing et al., 2012) and degradation of the endocochlear potential (Gratton et al., 1997; Schulte and Schmiedt 1992).

Collecting single-trial ABR and CAP data allows for the examination of inter-trial PLV. Neural synchrony decreased for human CAP N1 (Figure 6) and for mouse ABR wave I (Figure 4, left column) at all frequencies, demonstrating an age-related loss of temporal fidelity in AN responses. Mice and humans showed a decrease in response synchrony from the AN to the brainstem, yet response amplitudes were relatively preserved. These results are broadly consistent with prior reports following acute AN injury in mice: central gain recovers responses to rudimentary acoustic features, while precise spike timing remains disrupted (Chambers et al., 2016). Inconsistent response timing may manifest as perceptual deficits, especially in difficult listening environments, in which the signal is mixed with additional sources of noise.

The preservation of response amplitudes in the aging auditory midbrain, relative to AN responses, is likely the result of age-related decreases in inhibitory activity and/or decreases in the expression of inhibitory markers in the auditory brainstem and midbrain, which have been demonstrated in humans (Sharma et al., 2014) and in animal models (Caspary et al., 2008, Caspary & Llano, 2018), including CBA/CaJ mice (Tang et al., 2014). In older CBA/CaJ mice, SGNs show less GABA_A_R (inhibitory) α1 subunit expression and more NMDAR (excitatory) NR1 subunit expression, demonstrating an alteration of the balance of excitation and inhibition in the earliest stages of the auditory pathway (Tang et al., 2014). Furthermore, older CBA/J mice show significantly decreased glycine-mediated inhibition in the cochlear nucleus, which corresponds to increased firing rates in cochlear nucleus neurons (Frisina & Walton, 2006). Farther along the auditory pathway, in the inferior colliculus, single unit recordings reveal an age-related decrease in response latencies to amplitude modulated sounds in CBA/CaJ mice, which likely results from a shift in the excitation/inhibition (E/I) balance towards greater excitation (Simon et al., 2004). In summary, decreased afferent signaling leads to larger response amplitudes at higher auditory centers. Decreased neural synchrony at the level of the AN may be associated with a loss of cochlear synapses or changes in myelin structures. Deficits in neural synchrony may propagate through the auditory system resulting in the decreased midbrain synchrony observed in the current study. However, because inhibition is important for precise signal timing (Cardin, 2018) a loss of inhibition may introduce jitter to the timing of neural activity, which would be reflected in temporally variable latencies across responses, compounding deficits in synchrony in the AN.

If age-related disinhibition is the primary cause of central gain, then we would expect to see decreased response synchrony with relatively preserved response magnitudes in the midbrain. The results reported from both mice and humans in this study support this assertion, suggesting that changes in inhibitory signaling play an important role in age-related central gain. Within-subject studies, comparing inhibitory markers to measures of central gain, will further elucidate the role of age-related disinhibition.

## 5. Conclusions

In summary, we have demonstrated central gain in mice and humans, and further shown that age-related central gain does not ameliorate neural synchrony deficits. Future studies in mice will examine the precise relationship between measurements of age-related central gain and different pathophysiological aspects of aging, including demyelination (Xing et al., 2012), immune response (Noble et al., 2019), cochlear synaptopathy (Wu et al., 2019; Wu et al., 2020),and markers of inhibition and excitation, to better understand which factors have the greatest impact on age-related changes to neural synchrony and central gain. This may provide clinically relevant insights for treatments of age-related hearing deficits, which can be tested in the preclinical mouse model. Lastly, this work will inform future translational studies exploring the implications of deficient midbrain neural synchrony for cortical responses and auditory perception.

## CRediT Author Statement

**Jeffrey A Rumschlag:** Conceptualization, Methodology, Software, Formal analysis, Resources, Writing – Original Draft, Writing – Review & Editing, Visualization.

**Carolyn M. McClaskey:** Software, Formal analysis, Data Curation, Writing – Review & Editing.

**James W. Dias:** Formal analysis, Writing – Review & Editing.

**Lilyana B. Kerouac:** Investigation

**Kenyaria V. Noble:** Investigation

**Clarisse Panganiban:** Investigation

**Hainan Lang:** Conceptualization, Methodology, Investigation, Resources, Writing – Review & Editing, Supervision, Project administration, Funding acquisition.

**Kelly C. Harris:** Conceptualization, Methodology, Investigation, Resources, Writing – Review & Editing, Supervision, Project administration, Funding acquisition.

## Acknowledgements

We thank the participants of our study. We thank Brendan J. Balken and Carlene Brandon for their assistance with data collection.

## References

Brewton, D. H., Kokash, J., Jimenez, O., Pena, E. R., & Razak, K. A. (2016). Age-related deterioration of perineuronal nets in the primary auditory cortex of mice. Frontiers in Aging Neuroscience, 8(NOV), 1– 14. https://doi.org/10.3389/fnagi.2016.00270

Cai, R., Montgomery, S. C., Graves, K. A., Caspary, D. M. & Cox, B. C. The FBN rat model of aging: investigation of ABR waveforms and ribbon synapse changes. Neurobiology of Aging 62, 53–63 (2018). https://doi.org/10.1016/j.neurobiolaging.2017.09.034

Cardin, J. A. (2018). Inhibitory interneurons regulate temporal precision and correlations in cortical circuits. Trends in Neurosciences, 41(10), 689–700. https://doi.org/10.1016/j.tins.2018.07.015

Caspary, D. M., Ling, L., Turner, J. G., & Hughes, L. F. (2008). Inhibitory neurotransmission, plasticity and aging in the mammalian central auditory system. Journal of Experimental Biology, 211(11), 1781–1791. https://doi.org/10.1242/jeb.013581

Caspary, D. M., & Llano, D. A. (2018). Aging processes in the subcortical auditory system. The Oxford Handbook of the Auditory Brainstem.

Chambers, A. R., Resnik, J., Yuan, Y., Whitton, J. P., Edge, A. S., Liberman, M. C., & Polley, D. B. (2016). Central Gain Restores Auditory Processing following Near-Complete Cochlear Denervation. Neuron, 89(4), 867–879. https://doi.org/10.1016/j.neuron.2015.12.041

Cisneros-Franco, J. M., Ouellet, L., Kamal, B., & de Villers-Sidani, E. (2018). A brain without brakes: Reduced inhibition is associated with enhanced but dysregulated plasticity in the aged rat auditory cortex. ENeuro, 5(4). https://doi.org/10.1523/ENEURO.0051-18.2018

Dias, J.W., McClaskey, C.M., Harris, K.C., 2018. Time-compressed speech identification is predicted by auditory neural processing, perceptuomotor speed, and executive functioning in younger and older listeners. J. Assoc. Res. Otolaryngol. https://doi.org/10.1007/s10162-018-00703-1

Delorme, A. & Makeig, S. (2004). EEGLAB: an open-source toolbox for analysis of single-trial EEG dynamics. Journal of Neuroscience Methods,134(1), 9–21. https://doi.org/10.1016/j.jneumeth.2003.10.009

Frisina, R. D., & Walton, J. P. (2006). Age-related structural and functional changes in the cochlear nucleus. Hearing Research, 216–217(1–2), 216–223. https://doi.org/10.1016/j.heares.2006.02.003

Gmehlin, D., Kreisel, S. H., Bachmann, S., Weisbrod, M., & Thomas, C. (2011). Age effects on preattentive and early attentive auditory processing of redundant stimuli: is sensory gating affected by physiological aging?. Journals of Gerontology Series A: Biomedical Sciences and Medical Sciences, 66(10), 1043–1053. https://doi.org/10.1093/gerona/glr067

Gratton, M. A., Smyth, B. J., Lam, C. F., Boettcher, F. A., & Schmiedt, R. A. (1997). Decline in the endocochlear potential corresponds to decreased Na,K-ATPase activity in the lateral wall of quiet-aged gerbils. Hearing Research, 108(1–2), 9–16. https://doi.org/10.1016/S0378-5955(97)00034-8

Grose, J. H., Buss, E., & Elmore, H. (2019). Age-Related Changes in the Auditory Brainstem Response and Suprathreshold Processing of Temporal and Spectral Modulation. Trends in Hearing, 23, 1–11. https://doi.org/10.1177/2331216519839615

Harris K. C., Ahlstrom J. B., Dias J. D., Kerouac L., McClaskey C. M., Dubno J. R., Eckert M. A. (2021). Neural presbyacusis in humans inferred from age-related differences in auditory nerve function and structure. Journal of Neuroscience, 41(50), 10293–10304. https://doi.org/10.1523/JNEUROSCI.1747-21.2021

Harris, K. C., Vaden, K. I., McClaskey, C. M., Dias, J. W., & Dubno, J. R. (2018). Complementary metrics of human auditory nerve function derived from compound action potentials. Journal of Neurophysiology, 119(3), 1019–1028. https://doi.org/10.1152/jn.00638.2017

Harris, K. C., Wilson, S., Eckert, M. A., & Dubno, J. R. (2012). Human evoked cortical activity to silent gaps in noise: Effects of age, attention, and cortical processing speed. Ear and Hearing, 33(3), 330–339. https://doi.org/10.1097/AUD.0b013e31823fb585

Lang H, Kilpatrick LA, Samuvel DJ, Krug EL, Goddard JC (2011). Sox2 up-regulation and glial cell proliferation following degeneration of spiral ganglion neurons in adult mouse inner ear. J Assoc Res Otolaryngol. 12: 151–171. http://doi.org/10.1007/s10162-010-0244-1

Lopez-Calderon, J., & Luck, S. J. (2014). ERPLAB: An open-source toolbox for the analysis of event-related potentials. Frontiers in human neuroscience, 8, 213. https://doi.org/10.3389/fnhum.2014.00213

McClaskey, C. M., Dias, J. W., Dubno, J. R., & Harris, K. C. (2018). Reliability of measures of N1 peak amplitude of the compound action potential in younger and older adults. Journal of Speech, Language, and Hearing Research, 61(9), 2422–2430. https://doi.org/10.1044/2018_JSLHR-H-18-0097

McClaskey, C. M., Panganiban, C. H., Noble, K. V., Dias, J. W., Lang, H., & Harris, K. C. (2020). A multi-metric approach to characterizing mouse peripheral auditory nerve function using the auditory brainstem response. Journal of Neuroscience Methods, 346(September), 108937. https://doi.org/10.1016/j.jneumeth.2020.108937

Mohan, A., Thalamuthu, A., Mather, K. A., Zhang, Y., Catts, V. S., Weickert, C. S., & Sachdev, P. S. (2018). Differential expression of synaptic and interneuron genes in the aging human prefrontal cortex. Neurobiology of aging, 70, 194–202. https://doi.org/10.1016/j.neurobiolaging.2018.06.011

Möhrle, D., Ni, K., Varakina, K., Bing, D., Lee, S. C., Zimmermann, U., Knipper, M., & Rüttiger, L. (2016). Loss of auditory sensitivity from inner hair cell synaptopathy can be centrally compensated in the young but not old brain. Neurobiology of Aging, 44, 173–184. https://doi.org/10.1016/j.neurobiolaging.2016.05.001

Noble, K. V., Liu, T., Matthews, L. J., Schulte, B. A., & Lang, H. (2019). Age-related changes in immune cells of the human cochlea. Frontiers in Neurology, 10(AUG), 1–13. https://doi.org/10.3389/fneur.2019.00895

Ouda, L., Druga, R., & Syka, J. (2008). Changes in parvalbumin immunoreactivity with aging in the central auditory system of the rat. Experimental Gerontology, 43(8), 782–789. https://doi.org/10.1016/j.exger.2008.04.001

Ouellet, L., & de Villers-Sidani, E. (2014). Trajectory of the main GABAergic interneuron populations from early development to old age in the rat primary auditory cortex. Frontiers in Neuroanatomy, 8(JUN), 1–15. https://doi.org/10.3389/fnana.2014.00040

Parthasarathy, A., & Kujawa, S. G. (2018). Synaptopathy in the Aging Cochlea: Characterizing Early-Neural Deficits in Auditory Temporal Envelope Processing. The Journal of Neuroscience, 38(32), 7108–7119. https://doi.org/10.1523/JNEUROSCI.3240-17.2018

Pfefferbaum, A., Ford, J.M., Roth, W.T.F., Hopkins, W., Kopell, B.S. (1979). Event-related potential changes in healthy aged females. Electroencephalogr. Clin. Neurophysiol. 46 (1), 81–86. https://doi.org/10.1016/0013-4694(79)90052-X

Price, D., Tyler, L.K., Neto Henriques, R., Campbell, K.L., Williams, N., Treder, M.S., Taylor, J.R., Cam, C.A.N., Henson, R.N.A. (2017). Age-related delay in visual and auditory evoked responses is mediated by white-and grey-matter differences. Nat. Commun. 8, 15671. https://doi.org/10.1038/ncomms15671

Psatta, D. M., & Matei, M. (1988). Age-dependent amplitude variation of brain-stem auditory evoked potentials. Electroencephalography and Clinical Neurophysiology/ Evoked Potentials, 71(1), 27–32. https://doi.org/10.1016/0168-5597(88)90016-0

Rogalla, M. M., & Hildebrandt, J. K. (2020). Aging but not age-related hearing loss dominates the decrease of parvalbumin immunoreactivity in the primary auditory cortex of mice. eNeuro, 7(3). https://doi.org/10.1523/ENEURO.0511-19.2020

Sand, T. (1991). BAEP amplitudes and amplitude ratios: relation to click polarity, rate, age and sex. Electroencephalography and Clinical Neurophysiology, 78(4), 291–296. https://doi.org/10.1016/0013-4694(91)90183-5

Schrode, K. M., Muniak, M. A., Kim, Y. H., & Lauer, A. M. (2018). Central compensation in auditory brainstem after damaging noise exposure. eNeuro, 5(4). https://doi.org/10.1523/ENEURO.0250-18.2018

Schmiedt, R. A. (2010). The Physiology of Cochlear Presbycusis. 9–38. https://doi.org/10.1007/978-1-4419-0993-0_2

Schulte, B. A., & Schmiedt, R. A. (1992). Lateral wall Na, K-ATPase and endocochlear potentials decline with age in quiet-reared gerbils. Hearing Research, 61(1–2), 35–46. https://doi.org/10.1016/0378-5955(92)90034-K

Sergeyenko, Y., Lall, K., Liberman, M. C., & Kujawa, S. G. (2013). Age-Related Cochlear Synaptopathy: An Early-Onset Contributor to Auditory Functional Decline. Journal of Neuroscience, 33(34), 13686–13694. https://doi.org/10.1523/JNEUROSCI.1783-13.2013

Sharma, S., Nag, T. C., Thakar, A., Bhardwaj, D. N., & Roy, T. S. (2014). The aging human cochlear nucleus: Changes in the glial fibrillary acidic protein, intracellular calcium regulatory proteins, GABA neurotransmitter and cholinergic receptor. Journal of Chemical Neuroanatomy, 56, 1–12. https://doi.org/10.1016/j.jchemneu.2013.12.001

Simon, H., Frisina, R. D., & Walton, J. P. (2004). Age reduces response latency of mouse inferior colliculus neurons to AM sounds. The Journal of the Acoustical Society of America, 116(1), 469–477. https://doi.org/10.1121/1.1760796

Tang, X., Zhu, X., Ding, B., Walton, J. P., Frisina, R. D., & Su, J. (2014). Age-related hearing loss: GABA, nicotinic acetylcholine and NMDA receptor expression changes in spiral ganglion neurons of the mouse. Neuroscience, 259, 184–193. https://doi.org/10.1016/j.neuroscience.2013.11.058

Tremblay, K.L., Billings, C., Rohila, N., 2004. Speech evoked cortical potentials: effects of age and stimulus presentation rate. J. Am. Acad. Audiol. 15 (3), 226–237. https://doi.org/10.3766/jaaa.15.3.5

Ueno, H., Takao, K., Suemitsu, S., Murakami, S., Kitamura, N., Wani, K., Okamoto, M., Aoki, S., & Ishihara, T. (2018). Age-dependent and region-specific alteration of parvalbumin neurons and perineuronal nets in the mouse cerebral cortex. Neurochemistry International, 112, 59–70. https://doi.org/10.1016/j.neuint.2017.11.001

Xing, Y., Samuvel, D. J., Stevens, S. M., Dubno, J. R., Schulte, B. A., & Lang, H. (2012). Age-related changes of myelin basic protein in mouse and human auditory nerve. PLoS ONE, 7(4). https://doi.org/10.1371/journal.pone.0034500

Woods, D.L., Clayworth, C.C., 1986. Age-related changes in human middle latency auditory evoked potentials. Electroencephalogr. Clin. Neurophysiology Evoked Potentials Sect. 65 (4), 297–303. https://doi.org/10.1016/0168-5597(86)90008-0

Wu, P. Z., Liberman, L. D., Bennett, K., de Gruttola, V., O’Malley, J. T., & Liberman, M. C. (2019). Primary Neural Degeneration in the Human Cochlea: Evidence for Hidden Hearing Loss in the Aging Ear. Neuroscience, 407, 8–20. https://doi.org/10.1016/j.neuroscience.2018.07.053

Wu, P. Z., O’Malley, J. T., de Gruttola, V., & Charles Liberman, M. (2020). Age-related hearing loss is dominated by damage to inner ear sensory cells, not the cellular battery that powers them. Journal of Neuroscience, 40(33), 6357–6366. https://doi.org/10.1523/JNEUROSCI.093720.2020

